# Asynchronous viral spread of two unrelated viruses determines Lettuce Big Vein Disease symptom development

**DOI:** 10.1101/2025.02.13.638033

**Authors:** Willem E. W. Schravesande, Peter M. de Heer, Maurice Heilijgers, Adriaan Verhage, Harrold A. van den Burg

## Abstract

Lettuce big-vein disease (LBVD) is a major disease affecting lettuce cultivation worldwide. LBVD is caused by two unrelated negative-stranded RNA viruses, that is, Mirafiori lettuce big-vein virus (MiLBVV) and Lettuce big-vein associated virus (LBVaV) both vectored by the soilborne fungus *Olpidium virulentus*. Despite extensive research, a synergistic effect between the two viruses has not been observed, while both viruses individually have been suggested to be the causal agent for the disease. By performing lettuce reinfections using a large soil sample collection carrying LBVD infested *O. virulentus* spores, the presence of LBVaV was consistently established in diseased lettuce heads, while MiLBVV infections were apparently less prevalent. Yet, aboveground infections with MiLBVV corresponded with strong disease symptoms. Strikingly, the spread of LBVaV from the root to shoot always preceded that of MiLBVV. The LBVaV systemic spread was highly synchronized between plants, while MiLBVV spread was always delayed and asynchronous. A pan-genome analysis revealed independent segment reassortments for both viruses indicative of mixed field infections over the sampled period. Yet, RNA segment abundance was highly conserved for both viruses between all re-infections, suggesting that segment abundance has a regulatory role for the two individual viruses, but that segment abundance is not impacted by the presence of the other two viruses. The pan-genome analysis also revealed different evolutionary rates of the viral ORFs suggesting that mutagenesis of certain ORFs compromises viral fitness and thus revealing a potential weak spot for both viruses.

**Importance:** Lettuce big-vein disease (LBVD) is an important viral disease complex affecting lettuce cultivation worldwide. Here we reveal a synergistic interaction between the two principal associated segmented RNA viruses, Mirafiori lettuce big-vein virus (MiLBVV) and Lettuce big-vein associated virus (LBVaV). We show unequivocally that MiLBVV is the main virus responsible for severe disease symptoms in lettuce heads. Yet, MiLBVV root-to-shoot movement was in our conditions always preceded by LBVaV movement into the lettuce heads. Arguably, LBVaV thus facilitates the root-to-shoot movement of MiLBVV. Moreover, both viruses undergo segment reassortment increasing their genome plasticity and the reassortment events appeared to be independent events with mixed infections. Finally, we provide data that both viruses regulate gene expression via the copy number of their RNA segments, but that the genome formula does not change in dual infections. We thus provide evidence for a synergistic interaction needed for strong LBVD symptoms.

## Introduction

Lettuce big-vein disease (LBVD) is a disease complex affecting lettuce cultivation worldwide. Market value of plants suffering from this disease is diminished due to characteristic chlorosis along the veins, known as vein banding, coupled with stunted and distorted growth (1–3). Although LBVD was first described in 1934, a comprehensive pangenomic analysis of the viruses associated with the disease and their vector remains lacking. This knowledge gap has limited the depth of molecular characterization available for these viruses. LBVD is vectored by the soil-borne *Olpidium virulentus*, a chytrid fungus capable of producing persistent resting spores that remain viable for up to two decades in infected soils (4–6). This soil persistence presents a strong challenge for LBVD disease management. Its symptoms have been connected to the presence of at least two unrelated plant viruses, that is, Mirafiori lettuce big-vein virus (MiLBVV; *Ophiovirus mirafioriense*) and Lettuce big-vein associated virus (LBVaV; *Varicosavirus lactucae*). A recent report adds Lactuca big-vein associated phlebovirus (LBVaPV; *Olpivirus lactucae*) as a third virus associated with LBVD, although its role in LBVD remains to be solved (7, 8). LBVaV was the first virus characterized in LBVD-affected plants (9). Following its discovery, Koch’s postulates were seemingly fulfilled for this virus in 1987 (10). However, in 2000, MiLBVV was also found to be associated with LBVD (11) resulting in a need to re-address the causality question of the disease.

In fact, symptomatic plants commonly exhibit co-infections of LBVaV and MiLBVV (12–16). Nevertheless, studies using Enzyme-Linked Immunosorbent Assay (ELISA) and western-blotting to assess viral presence revealed earlier that lettuce plants solely infected with LBVaV did not exhibit LBVD symptoms (5, 17). In contrast, plants infected with MiLBVV showed typical symptoms, even in absence of LBVaV (13). Furthermore, no direct correlation was observed between the systemic spread of LBVaV and the development of LBVD symptoms. A notable correlation between symptom development and *in planta* spread of MiLBVV further stages this virus as the causal agent.

In contrast, Roggero *et al.* found no absolute correlation between the occurrence of LBVD symptoms and the presence of either of the two associated viruses and even reported two symptomatic plants infected by LBVaV but not by MiLBVV (12). Similarly, studies based on RT-qPCR detection of viral infections showed that symptomatic plants were present in the assays that only were tested positive for the presence of LBVaV and not for the presence of MiLBVV (13, 18–20). These discrepancies underscore the ambiguity about the precise roles of LBVaV and MiLBVV in LBVD pathology. An issue for resolving these inconsistencies is the limited genomic information available for the viruses implicated in LBVD: LBVaV, MiLBVV, and the recently described LBVaPV. The limited availability of genomic data has hampered the development of robust diagnostic tools and hindered our understanding of this disease complex. This scarcity in genomic data may also reduce the reliability of detection methods reported in earlier studies, calling for additional investigation.

The LBVaV genome consists out of two negative-sense, single-stranded RNA segments, approximately 6.8 kb (RNA 1) and 6.1 kb (RNA 2) in length. The virus belongs to the *Rhabdoviridae* family, genus varicosavirus (9). While its genome codes for at least six viral proteins, the function of four of its ORFs remains unclear (21–23). The MiLBVV genome consists of four negative-sense single-stranded RNA segments, approximately 7.8 kb (RNA 1), 1.7 kb (RNA 2), 1.5 kb (RNA 3), and 1.4 kb (RNA 4) in length. MiLBVV belongs to the *Ophioviridae* family, genus ophiovirus. The protein function of four of the seven viral ORFs is unknown (24–27). LBVaPV, unknown till recently, possesses a genome consisting of four RNA segments, approximately 6.4 kb (RNA 1), 1.1 kb (RNA 2), 1.4 kb (RNA 3), and 1.3 kb (RNA 4) in length. LBVaPV belongs to the *Phenuiviridae* family (7).

This study investigates the interaction between LBVaV and MiLBVV in LBVD, assessing their individual roles on symptom development, synergistic interactions and the systemic infection into the lettuce heads. By analyzing genomic data from diverse isolates, we examined viral co-occurrence patterns, RNA segment abundance, RNA segment reassortment, and the selection pressure on the ORFs coded for by these viruses. To further characterize their individual infection cycle dynamics, we tracked the temporal and spatial movement of the viruses in a time-course experiment. Together, our findings provide a new understanding on a potential synergistic interaction between MiLBVV and LBVaV on LBVD development and deepen our understanding of the interplay and individual roles of the two viruses for disease development.

## Results

### MiLBVV relies on LBVaV for the successful establishment of infection

To build a comprehensive genome view of the lettuce virome in cultivation areas affected by LBVD, a shotgun metagenomic sequencing approach was used. Thereto, a diverse set of 54 soil samples retrieved from LBVD-associated soils collected in different geographical locations and years, was used to re-infect plants in controlled experimental conditions. The lettuce virome was obtained using Oxford Nanopore (ONT) sequencing using total RNA extracted from plants testing positive for MiLBVV and/or LBVaV. In total, 34 different soil samples yielded one or more viral genomes (Figure 1A, Supplemental Table 1, Supplemental Table 2). The number of infections in the re-infected plants differed notably for both viruses, that is, reinfections with LBVaV exhibited a higher prevalence than reinfections with MiLBVV across our collection (Figure 1B). In fact, single infections of LBVaV were observed in nine occasions, while not a single case of a single infection of MiLBVV was found (Figure 1C). This observation suggests a potential role of LBVaV in facilitating MiLBVV infectivity and/or the MiLBVV re-infection could be more difficult from the stored soil samples suggesting that MiLBVV could be less persistent in our storage conditions of the soil samples. Moreover, a novel virus was identified in eight samples, for which we proposed the name Lactuca big-vein associated phlebovirus (LBVaPV; *Olpivirus lactucae*) (7). LBVaPV was consistently found together with both LBVaV and MiLBVV in these samples. This co-occurrence suggests a potential dependency among these viruses and/or it could indicate an important role for *O. virulentus* as a vector for a diverse set of plant viruses. The exact role of LBVaPV in LBVD pathology remains a subject of future studies.

**Figure 1.**
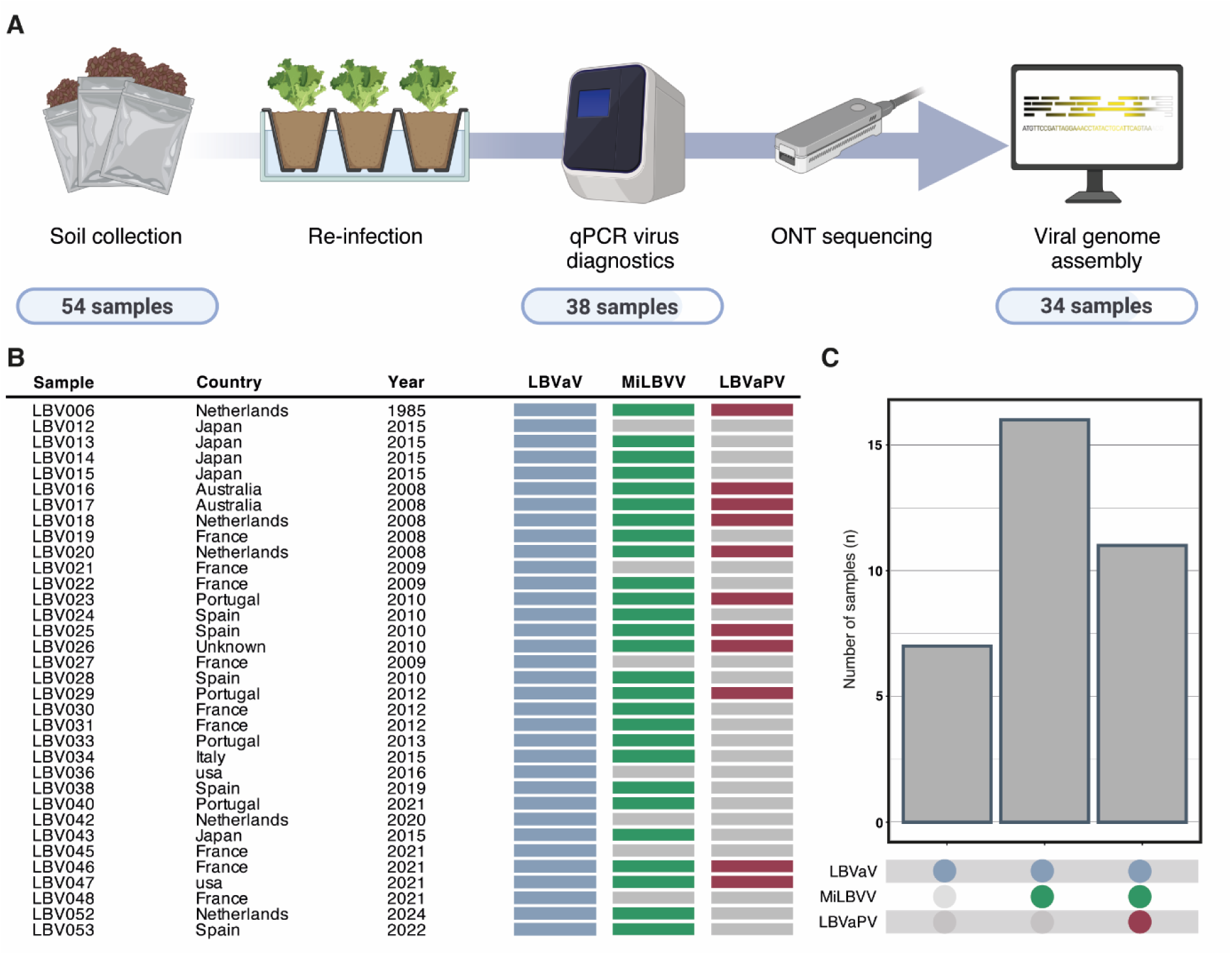
Presence of LBVaV is tightly connected with LBVD field infections. (A) Schematic representation of the experimental setup and the number of soil samples that passed each step in the workflow. (B) Overview of the samples with a complete viral genome assembly, including country of origin, year of collection, and presence/absence of each virus. (C) Visual overview of the number of reinfections yielding one or multiple viruses.

### MiLBVV and LBVaV facilitate segment reassortment, and these segments accumulate at specific frequencies

Segmentation of viral genomes, where the viral genome is divided into distinct, independently replicating segments, allows for efficient exchange of genetic information via segment reassortment within coinfected cells (28). These reassortments lead to novel combinations of genetic material that may promote virulence or allow host jump thereby enhancing viral plasticity. Using our soil sample collection, we assessed if we could find evidence for segment reassortment under field conditions for MiLBVV and LBVaV. Thereto, we performed a phylogenetic analysis at (a) the whole genome and (b) segment level to reveal segment reassortment for both viruses. Individual maximum likelihood (ML) trees were assembled for all viral segments using the complete coding region available in the genome, identifying clades within these trees (Supplemental Figure 1). When different RNA segments of a specific viral isolate do not group with the same clade, it provides strong evidence of segment reassortment. Several instances were observed where segments from the same isolate clustered with different phylogenetic clades, suggesting reassortment events for both MiLBVV and LBVaV (Figure 2A, B, indicated with black outlines). For example, RNA1 and RNA2 of the LBVaV isolate originating from soil sample LBV038 grouped with different viral clades, meaning that LBVaV-LBV038 has undergone reassortment (Supplemental Figure 1). Moreover, the individual viral segments of both viruses showed a systematic difference in sequence coverage depth. In fact, we observed a highly consistent pattern of relative viral segments abundances across the isolates collection, independent of a second virus being present or not (Figure 2C). Apparently, viral replication and/or viral transcription of the different RNA segments is highly controlled and confers some level of regulation on the two viruses in leaves.

**Figure 2.**
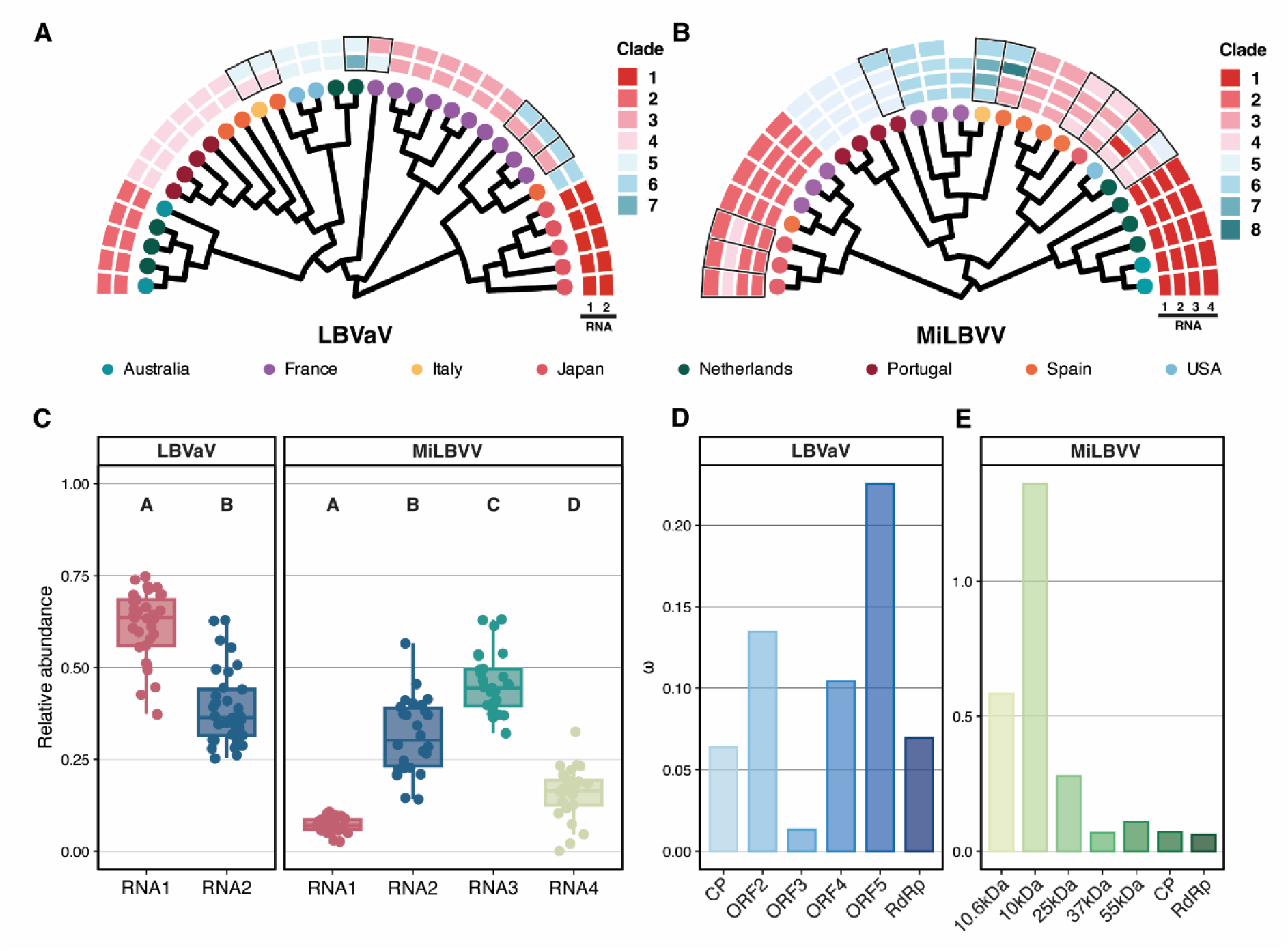
Phylogenetic analysis reveals independent segment reassortment for LBVaV and MiLBVV. (A,B) Phylogenetic tree of LBVaV (A) and MiLBVV (B), based on the concatenated genome, with tip node colors indicating the country of origin for each isolate. Additionally, for each isolate, individual RNA segments are depicted (outer rings), with metadata layers highlighting the clade to which each RNA fragment belongs (Supplemental figure 1). Isolates with indications of segment reassortment are marked with a black box. (C) Boxplot depicting the relative abundance of the RNA segments of LBVaV and MiLBVV. Each boxplot represents the distribution of relative abundance values for individual RNA segments across the two viruses (LBVaV: paired-t-test; p < 0.05 / MiLBVV: paired-t-test; p < 0.05). (D,E) Gene wide ω-ratio for the viral ORFs of LBVaV (D) and MiLBVV (E).

We also assessed the observed selection pressure on the viral proteins encoded by the different open reading frames (ORFs) by calculating the ratio of non-synonymous (dN) to synonymous substitutions (dS), also called ω ratio. This ratio reveals the degree and type of selection pressure on the different viral protein sequences, i.e. diversifying or purifying selection. Using the 34 LBVaV and 27 MiLBVV genomes, we performed a Branch-site Unrestricted Statistical Test for Episodic Diversification (BUSTED) (29) for each protein- coding ORF. For LBVaV, most ORFs exhibited a relative minor variation in the selection pressure. However, ORF3 displayed strong purifying selection, indicating that this viral protein tolerates little sequence variation and its function is important for viral fitness (Figure 2D). In contrast, MiLBVV showed distinct differences in selection pressure between the ORFs on RNA 4 and the other ORFs, indicating a diversifying selection pressure acts on the proteins encoded by RNA 4 (Figure 2E).

### MiLBVV root-to-shoot migration corresponds with Lettuce Big Vein Disease symptom development

To study the individual contributions of MiLBVV and LBVaV to symptom development, re-infections were performed with root material from a subset of re-infected plants showing lettuce head infections with (i) all three viruses, (ii) a double infection with MiLBVV and LBVaV, or (iii) only LBVaV alone(4 re-isolates per class, in total 12 isolates). Selected isolates varied in their origin and collection dates, ensuring a diverse set of virus samples, which enables the identification of potential isolate effects. Per strain, eight individual plants were inoculated, and disease symptom severity and viral titers were quantified nine weeks later. The individual viral titers were measured using qPCR, while the disease severity was scored using both a routine disease index score (DI-scores) and unbiased quantification of the relative chlorosis using an image analysis pipeline based on Ilastik (30), an interactive ML learning and segmentation toolkit (Figure 3A).

**Figure 3.**
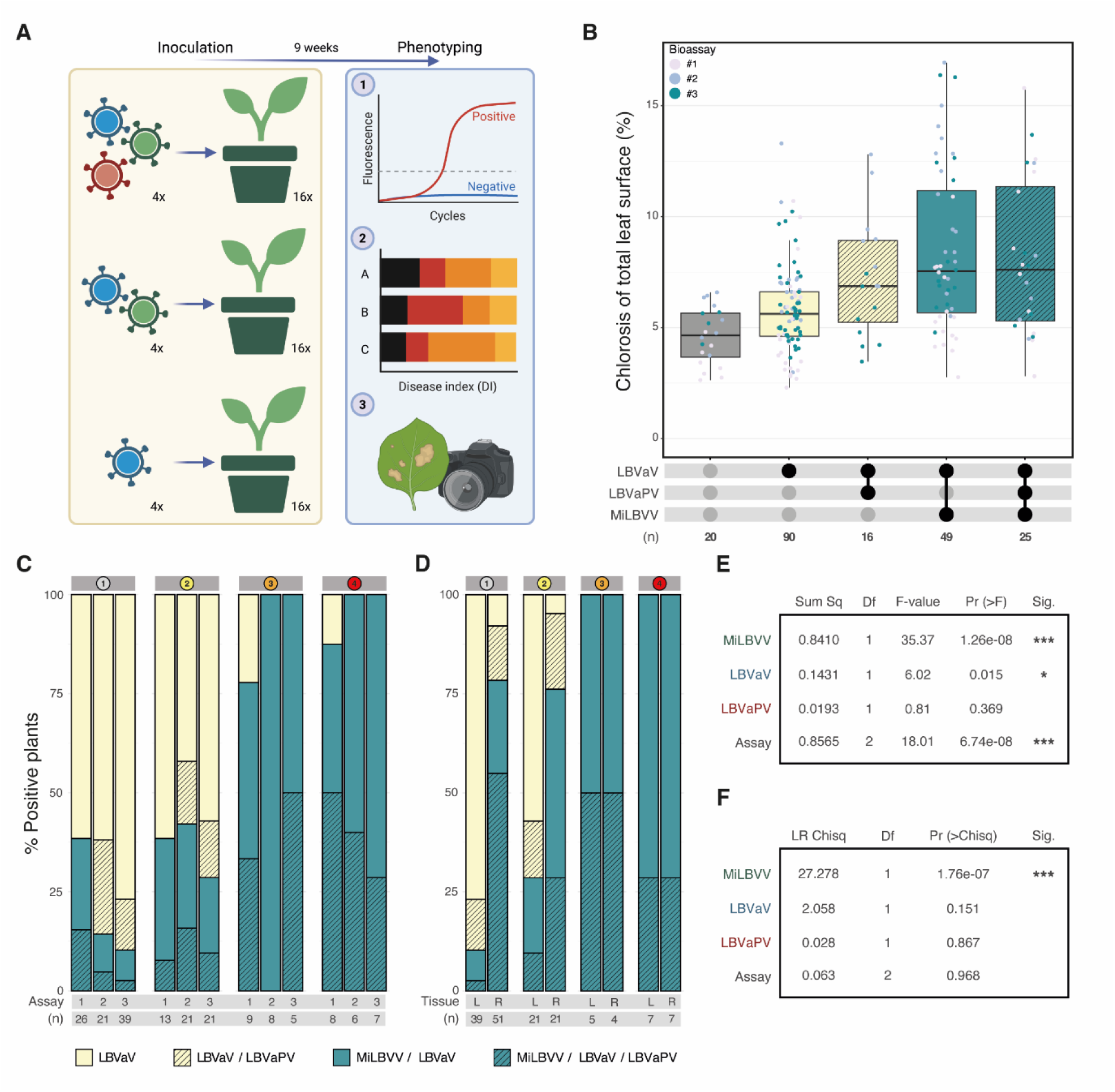
MiLBVV root-to-shoot migration is closely associated with LBVD symptom severity (disease index score of 3-4). (A) Schematic representation of the experimental set-up. Four isolates were selected per virus combination. Disease severity was assessed 9 weeks post inoculation using three methods, i.e., RT-qPCR, disease index (DI) scores and automated image analysis. (B) Disease symptom assessment quantified by the proportion of chlorotic leaf area (%). Presence/absence of each individual virus in individual leaves was assessed using RT-qPCR. Individual data point colors represent independent assays. (C) Stacked-bar plots facetted on DI-score; top bar 1=low symptoms to 4=high symptoms, showing the viral distribution over the DI scores for three independent repeats. (D) Viral distribution per tissue type, faceted by DI score. (E) Results of factorial ANOVA on the relative chlorotic leaf area and (F) DI.

As expected, the leaf chlorotic areas differed between plants with different viral combinations, but not within one combination. Plants infected with MiLBVV exhibited higher chlorosis levels compared to those containing only LBVaV or a combination of LBVaV with LBVaPV (Figure 3B). This observation was also reflected by the manually scored DI, indicating a positive correlation between presence of MiLBVV and LBVD symptom severity (Figure 3C). Using a factorial ANOVA we confirmed that MiLBVV is the main contributor to LBVD symptomology, that is, a major significant effect is seen for the presence of MiLBVV with leaf chlorosis development (F-value: 35.37, Pr(>F): 1.26e-08) with a minor effect for LBVaV (F-value: 6.02, Pr(>F): 0.015) (Figure 3E). For the DI-scores, the factorial ANOVA revealed only a significant effect for MiLBVV (LR Chisq: 27.2782, Df: 1, Pr(>Chisq): 1.762e-07), thus confirming that MiLBVV is the main contributor for strength of the LBVD phenotype (Figure 3F), although LBVaV alone can result in mild LBVD symptoms.

Next, we wondered if (a) the presence MiLBVV in the roots would already explain the disease symptom severity or (b) that spread of the two viruses to the above-ground tissue is needed for symptom development. Therefore, we sampled both leaf and root material from infected plants and determined again the viral titers in both plant parts (Figure 3D). Surprisingly, presence of MiLBVV in root tissue did not correlate with a high DI score. However, once MiLBVV had spread from the root to the shoot, symptom severity (and connected the DI score) increased. We thus conclude that MiLBVV root-to-shoot migration is closely associated with the sharp increase in LBVD symptoms (DI 3-4).

### MiLBVV and LBVaV exhibit distinct temporal and spatial patterns

To expose in more detail the dynamics of the interaction between MiLBVV, LBVaV, and LBVaPV during viral replication and systemic movement, a comprehensive timeseries experiment was conducted over a period of 64 days. The viral titer of each virus was determined during this period, in both root and shoot material (Figure 4). In root tissues, the titers of MiLBVV and LBVaV exhibited a rapid but synchronous increase during the early stages of infection, reaching a plateau level at approximately 15–20 dpi. The uniformity observed in these first stages of the infection is in sharp contrast with the dynamic pattern observed in the shoots. LBVaV consistently initiated replication in the shoot prior to MiLBVV with a systemic spread that started at approximately 30 dpi and reached a plateau by 40 dpi. In contrast, the systemic spread of MiLBVV to the aboveground tissues appeared to be highly stochastic, with a remarkable variation in the spread between plants both in timing and magnitude of shoot colonization. Interestingly, the more rapid and more synchronous movement of LBVaVs from the root-to-shoot was consistently seen for prior to MiLBVV spread/replication in the aboveground tissues. This observation aligns with our earlier observation that MiLBVV (Figure 1) was never detected in the shoots of plants where LBVaV was absent. MiLBVV exhibited delayed and highly variable systemic movement, with the root-shoot jump being a stochastic event for individual plants. Notably, LBVaPV, despite its presence in roots, failed to migrate systemically in this experiment, reinforcing its distinct role—or potential lack thereof—in the LBVD complex under our test conditions. These findings highlight the distinct spatiotemporal infection strategies employed by LBVaV and MiLBVV and reveal a potential interdependence in systemic spread and disease progression.

**Figure 4.**
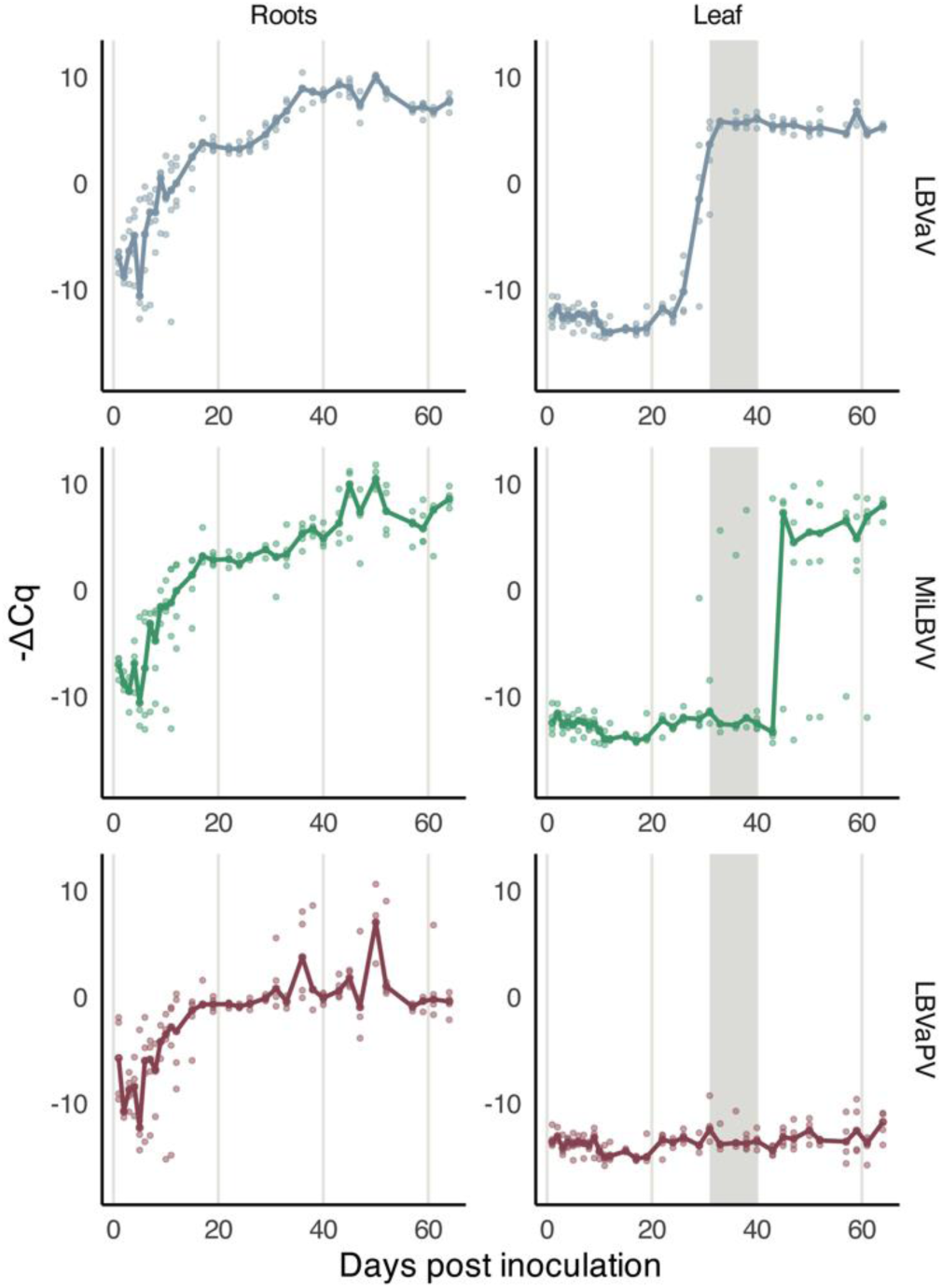
Time-course experiment revealing the asynchronous spread of LBVaV and MiLBVV, and LBVaPV) from the lettuce root to shoot. Over a period of 64 days, root and shoot material was sampled at 33 individual timepoints (4 replicates). Viral titers were quantified using RT-qPCR. The solid lines represent the median viral titers across four biological replicates. The shaded region indicates the time window where significant differences in the viral spread were observed between MiLBVV and LBVaV (ANOVA; p < 0.05).

## Discussion

Despite extensive research on LBVD, the precise role of the two viruses MiLBVV and LBVaV in disease symptomology remains disputed. We here resolved the lack of comprehensive genomic data for both viruses and generated a more holistic view on LBVD symptom development evaluating the role of the two viruses jointly.

We confirm that LBVD arises from a complex interaction between LBVaV and MiLBVV, with (i) MiLBVV being the primary contributor to symptom development and (ii) LBVaV apparently playing a crucial supporting role for MiLBVV replication and/or systemic spread. Our findings suggest that disease resistance against LBVaV could also present an agronomic solution for LBVD complex in lettuce cultivars, as a single resistance gene against LBVaV could suffice to suppress the root-to-shoot migration of MiLBVV and thereby giving symptom free plants. Likely, the MiLBVV viral titers would remain high in lettuce roots of such LBVaV- resistant plants, thus such a resistance gene would provide *in trans* some level of tolerance to a MiLBVV (root) infection (31). It would be of interest to examine if such a genetic source of LBVaV resistance would also suppress vector transmission by *O. virulentus* although one would need a virus-free culture of *O. virulentus* for this experiment. The consistent joint presence of these viruses in lettuce heads with severe LBVD symptoms suggests at least a synergistic or beneficial interaction for MiLBVV in disease onset and/or development. We reveal two phases in the infection, where LBVaV consistently establishes its root-to-shoot movement prior to that of MiLBVV. This could indicate a beneficial host manipulation by LBVaV that than supports the spread of MiLBVV. Moreover, our pangenome analysis revealed evidence of viral segment reassortment and conserved segment abundance across isolates, suggesting tight regulatory control of the viral segment replication.

These results support the notion that MiLBVV is closely associated with LBVD symptoms, while LBVaV alone does not induce strong symptoms, as also suggested by Sasaya *et al.* (17). However, the results from earlier studies investigating the roles of LBVaV and MiLBVV in LBVD remained inconclusive, possibly due to differences in diagnostic tests. Roggero et al. used ELISA for detecting LBVaV and MiLBVV, which is less sensitive than RT-PCR-based detection assays (12). This might have led to missed infections (false negatives) and inconsistent findings regarding virus presence and symptoms. Alemzadeh *et al.* and Navarro et al. used PCR-based assays, but we noted that the lack of high-quality genomic data might have led to unreliable primer design, resulting in false negatives and a lowered sensitivity (13, 18)(Supplemental figure 2). Hernandez et al. used the same primers thus facing similar challenges (19). These limitations could explain the discrepancies between the different earlier studies and highlight a need for more robust diagnostic tools. Our pangenome data allows at least for an optimal and robust design of primer pairs for both viruses.

Furthermore, the evidence of segment reassortment in MiLBVV and LBVaV aligns with previous reports on the reassortment capacity of multipartite and segmented viruses (28). By now it is evident that exchange of genetic viral segments contributes to genome plasticity and thereby viral fitness of the isolates. Reassortment between different tospoviruses has been shown to result in novel genomic combinations (32), which may occur in a similar fashion in MiLBVV and LBVaV, enhancing their plasticity and fitness in different environments. This ability to exchange genome segments serves as an evolutionary mechanism that provides RNA viruses with enhanced genome plasticity, potentially aiding their adaptation to new hosts or overcoming plant resistance mechanisms (33).

The concept of a stable genome copy formula highlights the importance of segment abundance in regulating expression of viral ORFs and maintaining viral fitness (34). For the nanovirus (a circular single-stranded DNA virus) Faba bean necrotic stunt virus (FBNSV) and the multipartite RNA virus Beet yellow vein virus (BNYVV) genomic segment abundance was shown to regulate gene expression, with FBNSV segment ratios also adapting to changes in the host environment to ensure optimal viral fitness and replication (35–37). Differences in segment copy number have been shown to directly influence gene expression, highlighting the plasticity of multipartite viruses in response to different conditions. Our data suggests that both MiLBVV and LBVaV may employ a similar strategy to regulate their protein abundances, where segment ratios are tightly controlled to optimize viral replication and spread. However, we found no impact of the presence of MiLBVV on the LBVaV genome copy formula (GCN). It will be interesting to see if the GCN differs between the root and shoot for either virus. Currently, this question cannot be answered as our root tissue samples might also capture (in part) viral replication in the vector *O. virulentus.* Hence, we would have to develop an *O. virulentus-*free viral infection, e.g., using *in vitro* tissue grafting, mechanical inoculation or inoculation with infectious clones. In MiLBVV and LBVaV, such genome copy regulation could help maintain an effective balance of the viral proteins necessary for a successful infection and systemic movement.

An intriguing question remains regarding the manner in which LBVaV facilitates MiLBVV replication and systemic spread. Despite the wealth of evidence on different synergistic and antagonistic interactions when different viruses (attempt to) infect the same host plant, simultaneously or in a sequential order, the biological significance of these interactions remains extremely difficult to prove in absence of infectious clones or single infections (38, 39). One plausible explanation is that LBVaV may function as a helper virus in the LBVD complex. Helper-dependent viruses rely on co-infecting “helper” viruses to provide viral functions *in trans*, which they cannot efficiently execute on their own (40, 41). This phenomenon is well documented in plant virus systems; for instance, umbraviruses depend on co-infecting luteovirids for encapsidation and vector transmission (42, 43). In our study, LBVaV’s consistent establishment of root-to-shoot movement ahead of MiLBVV raises the possibility that LBVaV complements functions—perhaps in viral movement or in suppression of host defenses—that MiLBVV alone cannot perform. Such a helper-dependent interaction not only explains the synergistic relationship between the two viruses but also aligns with rare documented cases of viral interdependence in plant disease complexes, as seen in the mutualistic relationship observed in Pea enation mosaic virus (PEMV) (44, 45). Future research should focus on elucidating the molecular mechanisms by which LBVaV facilitates MiLBVV infection, e.g. long-distance viral movement. Intriguingly, expression of the 54kDa protein encoded on the RNA2 genomic segment of MiLBVV resulted in intercellular movement of the movement-defective Tomato mosaic virus (ToMV) in *N. benthamiana* (26, 46, 47). This would mean that MiLBVV would at least be able to move from cell-to-cell via plasmodesmata, but it does not detail if this candidate movement protein of MiLBVV also promotes the long-distance phloem transport of plant viruses in the lettuce aboveground tissue. Investigating the specific host or fungal factors influenced by LBVaV is thus likely to provide us with new insights on here proposed synergistic interaction between these viruses in terms of systemic spread and/or vector transmission. Additionally, more in-depth studies on LBVaPV are needed to determine its role, if any, in the progression of LBVD, particularly under different environmental conditions

In conclusion, our results reveal that MiLBVV is the primary contributor to LBVD symptoms, with LBVaV playing an apparent key role in facilitating systemic infection of MiLBVV, highlighting their interdependence and the complexity of LBVD pathogenesis.

## Material & Methods

### Plant material

*Lactuca sativa* cultivar ‘IGLO’ plants were sown on potting soil and grown at 14°C for 2 weeks. For the screening of the LBVD soil sample isolate collection, seedlings were transplanted on infected soil samples that originating from natural infections in lettuce cultivation. Soil samples were collected over multiple years from different counties of origin and stored in the dark at 4°C until use. A complete overview of the soil samples is provided in Supplemental Table 1. Eight to twelve weeks post inoculation, the presence of MiLBVV and LBVaV was assessed using Taqman RT-qPCR (Supplemental Table 3). In short, 100 mg of plant material (leaf, root) was sampled from each plant and used for total RNA isolation using the RNeasy Plant Mini Kit (Qiagen). Unless otherwise stated, manufacturers protocols were followed. RT-qPCR was performed using specific Taqman assays for MiLBVV, LBVaV, and the *Lactuca sativa* housekeeping gene TIP41 (primers are detailed in the Supplemental Table 3). Infected material, from plants that were positive for either MiLBVV and/or LBVaV, was directly flash frozen with use of liquid nitrogen for later processing.

For the disease assays, *L. sativa* cultivar ‘Green Towers’, plants were sown on potting soil and grown at 14°C for 2 weeks. *L. sativa* cultivar ‘Green Towers’ was selected due to its reduced susceptibility to *Botrytis cinerea*. A subset of the isolate collection was selected based on presence / absence of the viruses using the country of origin and collection year as a way to have a diverse panel of isolates. In total, 12 different isolates were selected for further analysis. Infected soil material was mixed in a 1:6 ratio with clean potting soil and manually homogenized to a uniform mixture. This soil mixture was used to fill pots 12 cm in diameter, in which 2-week-old seedlings were transplanted. To avoid cross contamination between the different isolates, the isolates were placed in individual watering trays with sufficient spacing between the different trays.

For the timeseries, *L. sativa* cultivar ‘Green Towers’ plants were sown on potting soil and pre-grown at 14°C for 2 weeks. The seedlings were then transplanted into a hydroponic gutter system, where *O. virulentus* spores containing the viruses of interest circulated due to the presence of infected plants, with one infected row of plants present for every ten rows of plants. At each individual timepoint, 4 plants were sampled. Roots were washed with a gentle hose to remove soil and debris, after which root and shoot were separated and directly flash frozen with use of liquid nitrogen for later processing.

### Whole genome sequencing

Total RNA was isolated from the flash-frozen infected *L. sativa* material. Unless otherwise stated, manufacturers protocols were followed. Material was grinded using mortar and pestle to a fine powder and used as input material for total RNA isolation using the RNeasy Plant Mini Kit (Qiagen). The isolated total RNA was subjected to ribosomal depletion using the RiboMinus Plant Kit for RNA-Seq (Thermo Scientific). The remaining RNA was used for cDNA synthesis using the Maxima H Minus Double Stranded cDNA Synthesis Kit using random hexamer primers (Thermo Scientific). Oxford Nanopore library construction was performed with ds-cDNA as input. For library construction, the Native Barcoding Kits SQK-NBD114.96 (Oxford Nanopore) was used. Manufacturer’s instructions and protocols were followed. The constructed sequencing libraries were sequenced using the Promethion FLO-PRO114M flowcells (Oxford Nanopore) during a period of 72 hours.

### Genome assembly and analysis

The sequencing data was processed on a high compute cluster. The raw sequencing data was basecalled using Dorado v0.5.0 (https://github.com/nanoporetech/dorado) followed by QC filtering using Nanofilt v2.2.0 (48) (https://github.com/wdecoster/nanofilt). The filtered data was subjected to barcode demultiplexing using qcat v1.1.0 (https://github.com/nanoporetech/qcat), splitting the multiplexed samples into individual bundle files. Sequencing adapters were computationally trimmed of using porechop (https://github.com/rrwick/Porechop). To filter out the plant genome present in the sequencing data, the data was mapped to the reference genome of *L. sativa* ‘Salinas’ (v14)(49) using minimap2 v2.16 (50) (https://github.com/lh3/minimap2). The unmapped reads were exported into a separate bundle file, that was used as input data for *de novo* genome assembly using Flye v2.9.3 (51) (https://github.com/fenderglass/Flye). The genome assemblies were polished using Medaka v1.11.3 (https://github.com/nanoporetech/medaka), followed by manual curation and annotation (Supplemental Table 2). The raw reads were deposited under BioProject accession number <u>PRJNA1211292</u>.

A phylogenetic overview of the sequenced genomes of MiLBVV and LBVaV was created using Geneious Prime 2022.x.x (https://www.geneious.com) in combination with R studio v2023.03.0. Analysis of presence and rates of (non-)synonymous SNPs was performed using HyPhy package (52) (https://www.hyphy.org), which is a package for comparative sequence analysis using evolutionary models. For this specific question, the BUSTED (Branch-site Unrestricted Statistical Test for Episodic Diversification) (29) analysis was used. For LBVaV, relative abundances of RNA1 and RNA2 were compared using a paired t-test. Significant differences were identified at a threshold of p<0.05. For MiLBVV, pairwise comparisons were conducted using paired t-tests. To account for multiple testing, p-values were adjusted using the Bonferroni and Benjamini-Hochberg (BH) correction methods, with adjusted p-values below p<0.05 considered significant.

### Disease assays and timeseries

Nine weeks post infection, the disease assays were scored and the individual plants sampled. A disease score was given to each plant using an ordinal scale ranging from 1 to 4 based on the observed symptoms per plant (Supplemental Table 4). Secondly, material was sampled from each plant for RT-qPCR analysis. Lastly, a representative leaf from similar developmental stages, was photographed for each plant.

Presence of MiLBVV, LBVaV and LBVaPV was assessed by RT-qPCR for disease assays and timeseries. In short, 100 mg of leaf material was sampled from each plant and used for total RNA isolation using the innuPREP RNA Virus PLUS kit (Analytic Jena) on the Kingfisher Flex platform (Thermofisher Scientific), following manufacturer’s instructions. RT-qPCR was performed using specific Taqman assays for MiLBVV, LBVaV, and the *L. sativa* housekeeping gene *TIP41* (Supplemental Table 3) using the Taqman Fast Virus 1-step Master Mix (Thermofisher Scientific). Pictures were analyzed using Ilastik, a toolkit for image segmentation and learning (30). The image was classified into 6 different categories, enabling the quantification of chlorosis symptoms associated with LBVD. After classification, pixel counts were generated with use of ImageJ for each category.

All statistical analyses were performed using R (v4.3.1) with packages including tidyverse (v2.0.0), lme4 (v.1.1-35.5), glmmTMB (v.1.1.9), car (v3.1-2), and performance (v.0.12.2). The distribution of variables was assessed, and correlation between response variables was computed using Spearman’s method. Various models were tested, including Gaussian, Inverse Gaussian, Gamma, Poisson, quasi-Poisson, negative binomial and Gaussian with log transformation, which were used in order to determine the best fit. The selected model (% chlorosis: normal distribution with log10 transformation; DI: Poisson distribution) was further evaluated using diagnostic plots and used to perform a factorial ANOVA.

---

**Supplemental Figure 1.**
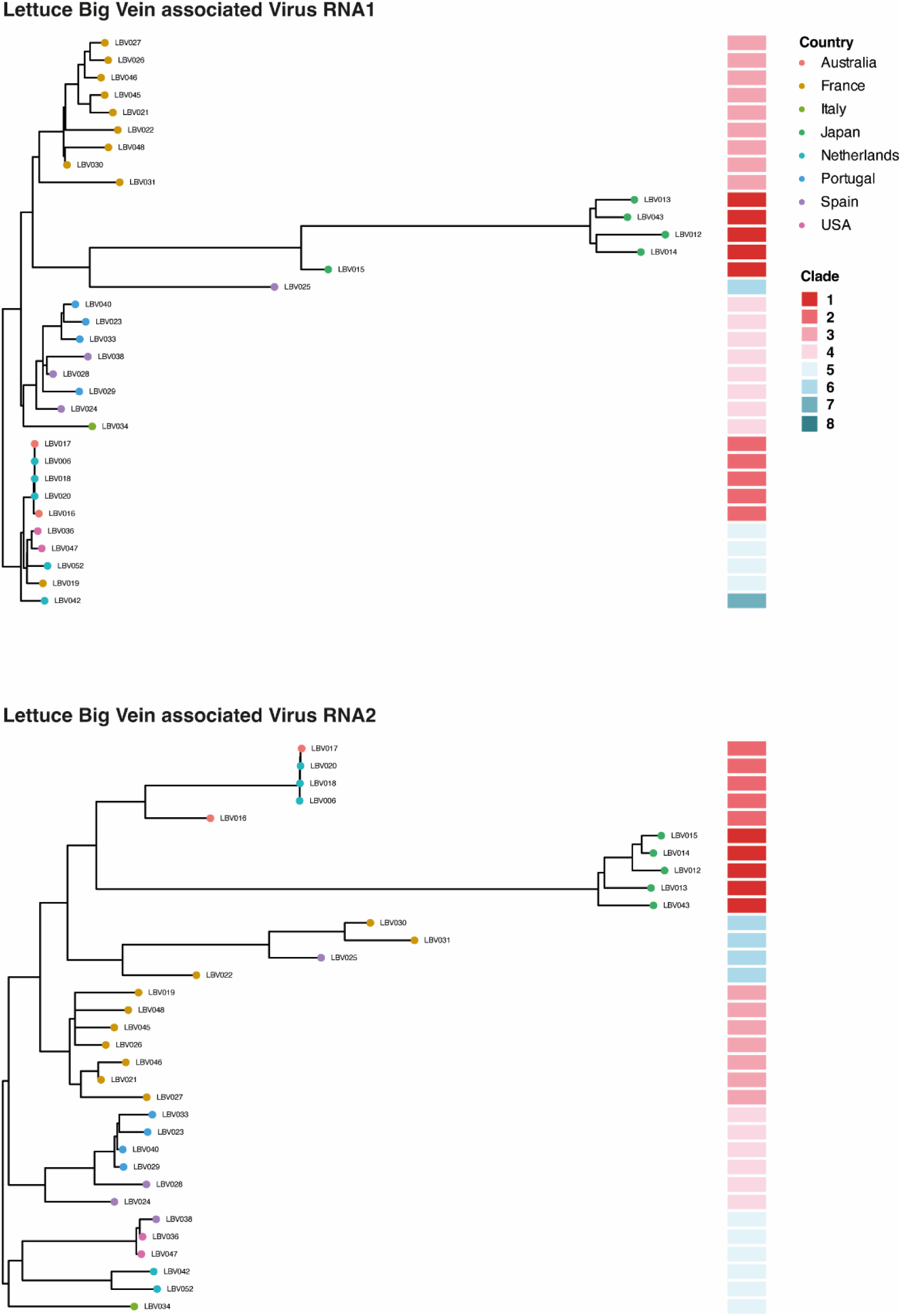
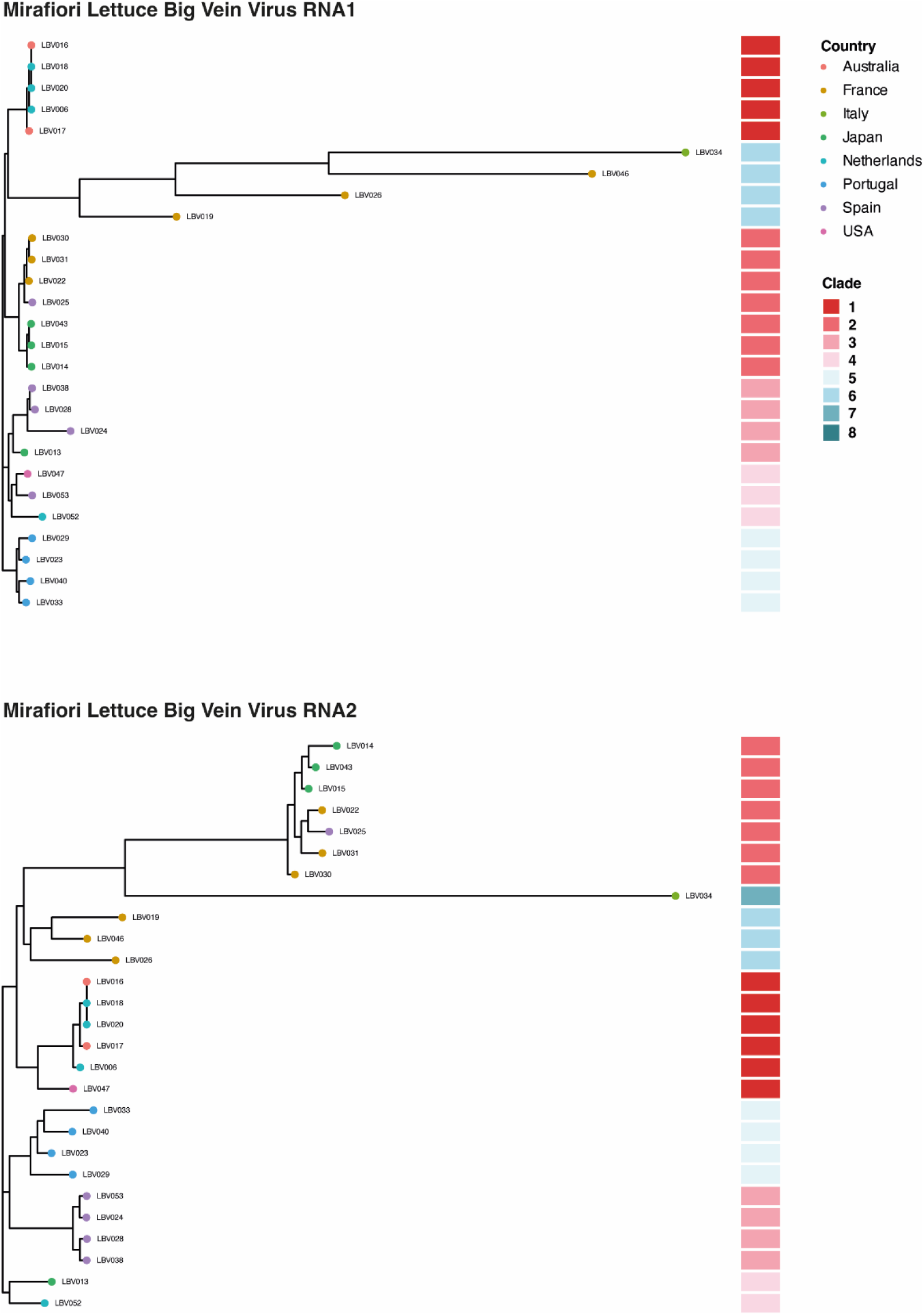
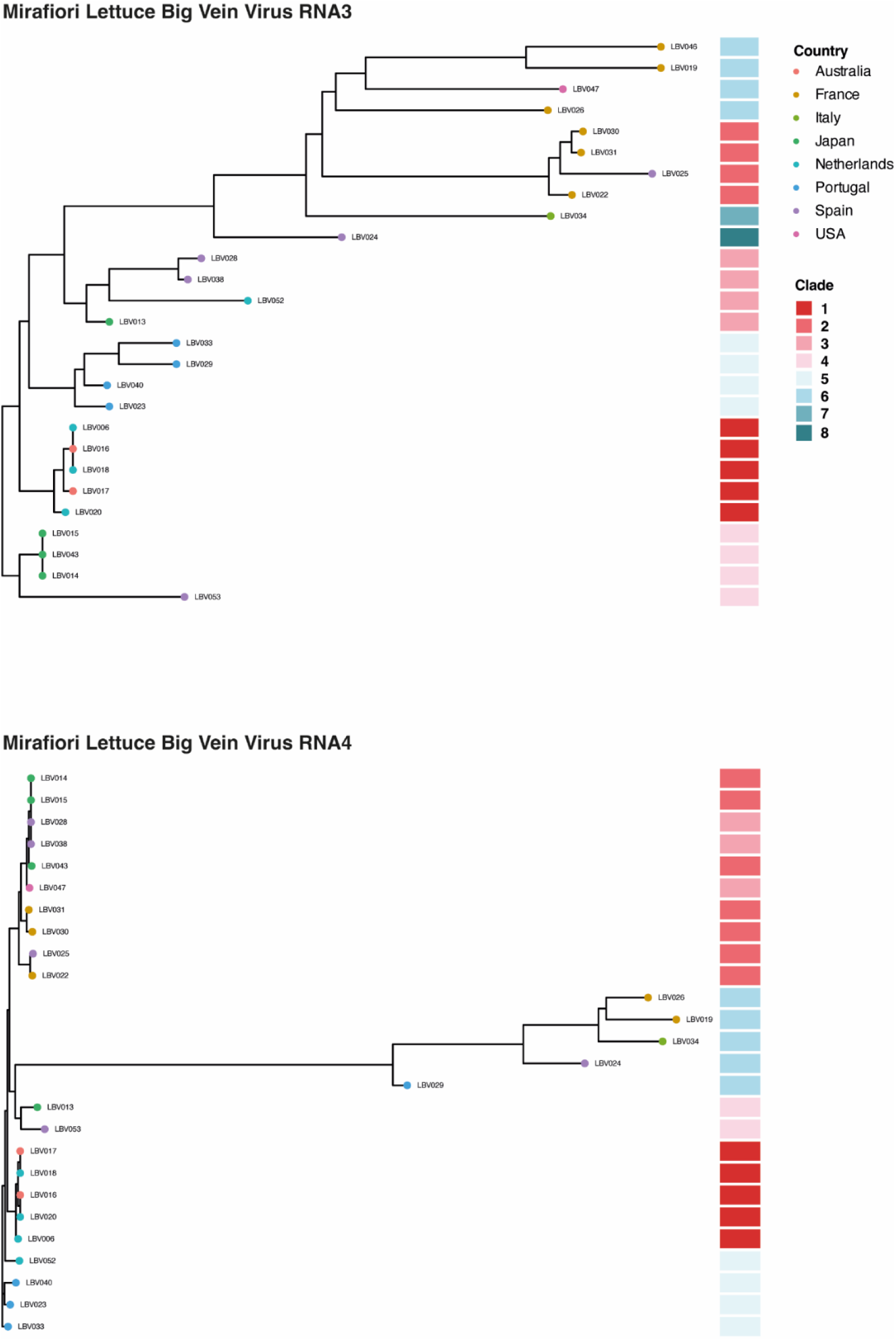
Phylogenetic Trees per RNA segment. Maximum likelihood (ML) phylogenetic trees for individual RNA segments of LBVaV and MiLBVV isolates, with node colors representing country of origin and color blocks representing the phylogenetic clades.

**Supplemental Figure 2.**
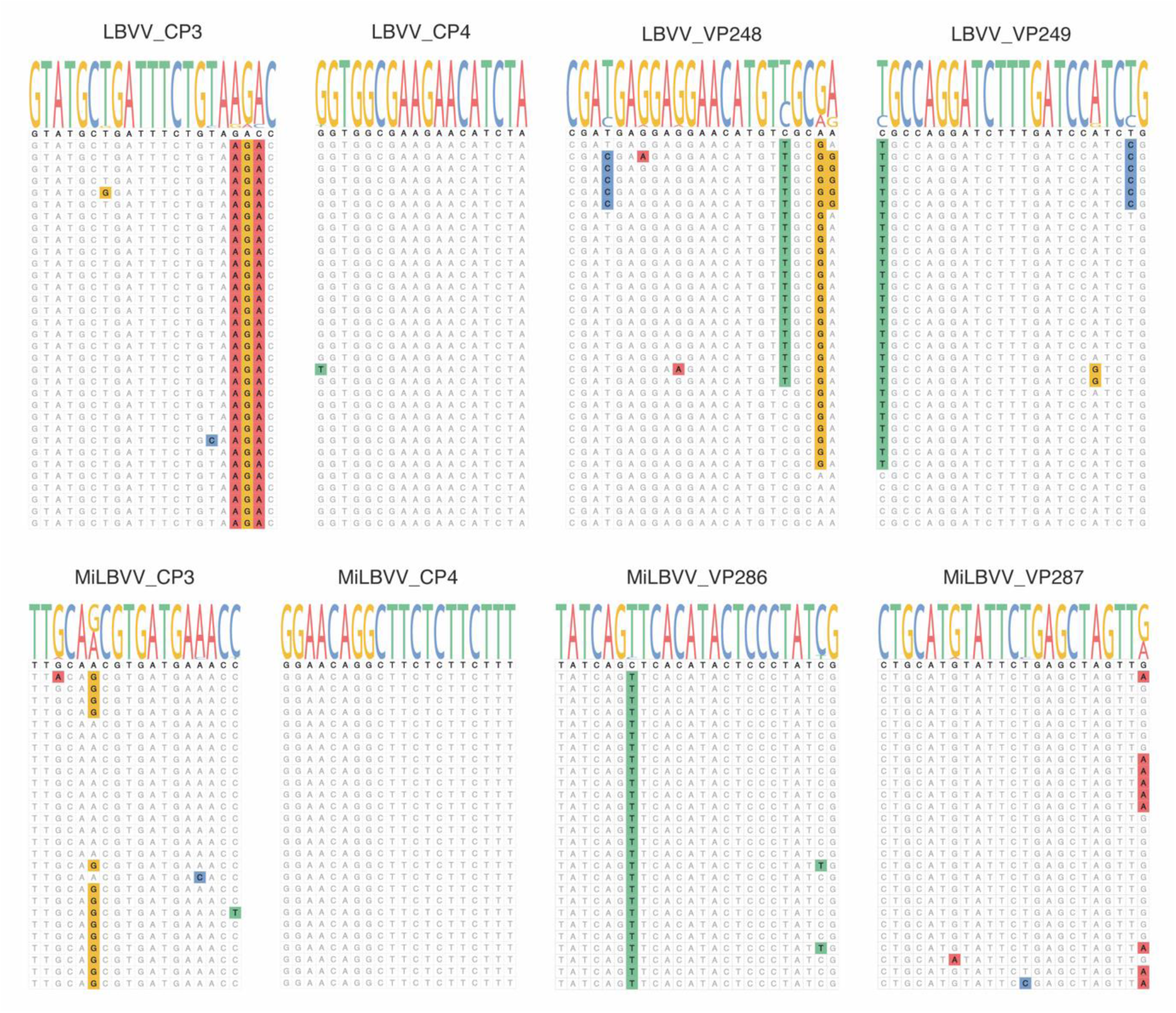
Legacy primer binding sites. Diagram illustrating the binding sites of primers used in diagnostic assays for detecting LBVaV and MiLBVV. Primers were mapped to their respective genomic locations, highlighting mismatches between the legacy primers and newly obtained viral genome sequences.

**Supplemental Table 1.** Overview of LBVD-associated soil samples. Table summarizing the isolates included in the study, with details such as sample ID, country of origin, host plant, and year of collection.

| Sample # | Country | Host | Year | Accession |
| --- | --- | --- | --- | --- |
| LBV006 | Netherlands | *Lactuca sativa* | 1985 | SAMN46272366 |
| LBV012 | Japan | *Lactuca sativa* | 2015 | SAMN46272367 |
| LBV013 | Japan | *Lactuca sativa* | 2015 | SAMN46272368 |
| LBV014 | Japan | *Lactuca sativa* | 2015 | SAMN46272369 |
| LBV015 | Japan | *Lactuca sativa* | 2015 | SAMN46272370 |
| LBV016 | Australia | *Lactuca sativa* | 2008 | SAMN46272371 |
| LBV017 | Australia | *Lactuca sativa* | 2008 | SAMN46272372 |
| LBV018 | Netherlands | *Lactuca sativa* | 2008 | SAMN46272373 |
| LBV019 | France | *Lactuca sativa* | 2008 | SAMN46272374 |
| LBV020 | Netherlands | *Lactuca sativa* | 2008 | SAMN46272375 |
| LBV021 | France | *Lactuca sativa* | 2009 | SAMN46272376 |
| LBV022 | France | *Lactuca sativa* | 2009 | SAMN46272377 |
| LBV023 | Portugal | *Lactuca sativa* | 2010 | SAMN46272378 |
| LBV024 | Spain | *Lactuca sativa* | 2010 | SAMN46272379 |
| LBV025 | Spain | *Lactuca sativa* | 2010 | SAMN46272380 |
| LBV026 | France | *Lactuca sativa* | 2010 | SAMN46272381 |
| LBV027 | France | *Lactuca sativa* | 2009 | SAMN46272382 |
| LBV028 | Spain | *Lactuca sativa* | 2010 | SAMN46272383 |
| LBV029 | Portugal | *Lactuca sativa* | 2012 | SAMN46272384 |
| LBV030 | France | *Lactuca sativa* | 2012 | SAMN46272385 |
| LBV031 | France | *Lactuca sativa* | 2012 | SAMN46272386 |
| LBV033 | Portugal | *Lactuca sativa* | 2013 | SAMN46272387 |
| LBV034 | Italy | *Lactuca sativa* | 2015 | SAMN46272388 |
| LBV036 | USA | *Lactuca sativa* | 2016 | SAMN46272389 |
| LBV038 | Spain | *Lactuca sativa* | 2019 | SAMN46272390 |
| LBV040 | Portugal | *Lactuca sativa* | 2021 | SAMN46272391 |
| LBV042 | Netherlands | *Lactuca sativa* | 2020 | SAMN46272392 |
| LBV043 | Japan | *Lactuca sativa* | 2015 | SAMN46272393 |
| LBV046 | France | *Lactuca sativa* | 2021 | SAMN46272394 |
| LBV047 | France | *Lactuca sativa* | 2021 | SAMN46272395 |
| LBV052 | France | *Lactuca sativa* | 2021 | SAMN46272396 |

**Supplemental Table 2.**
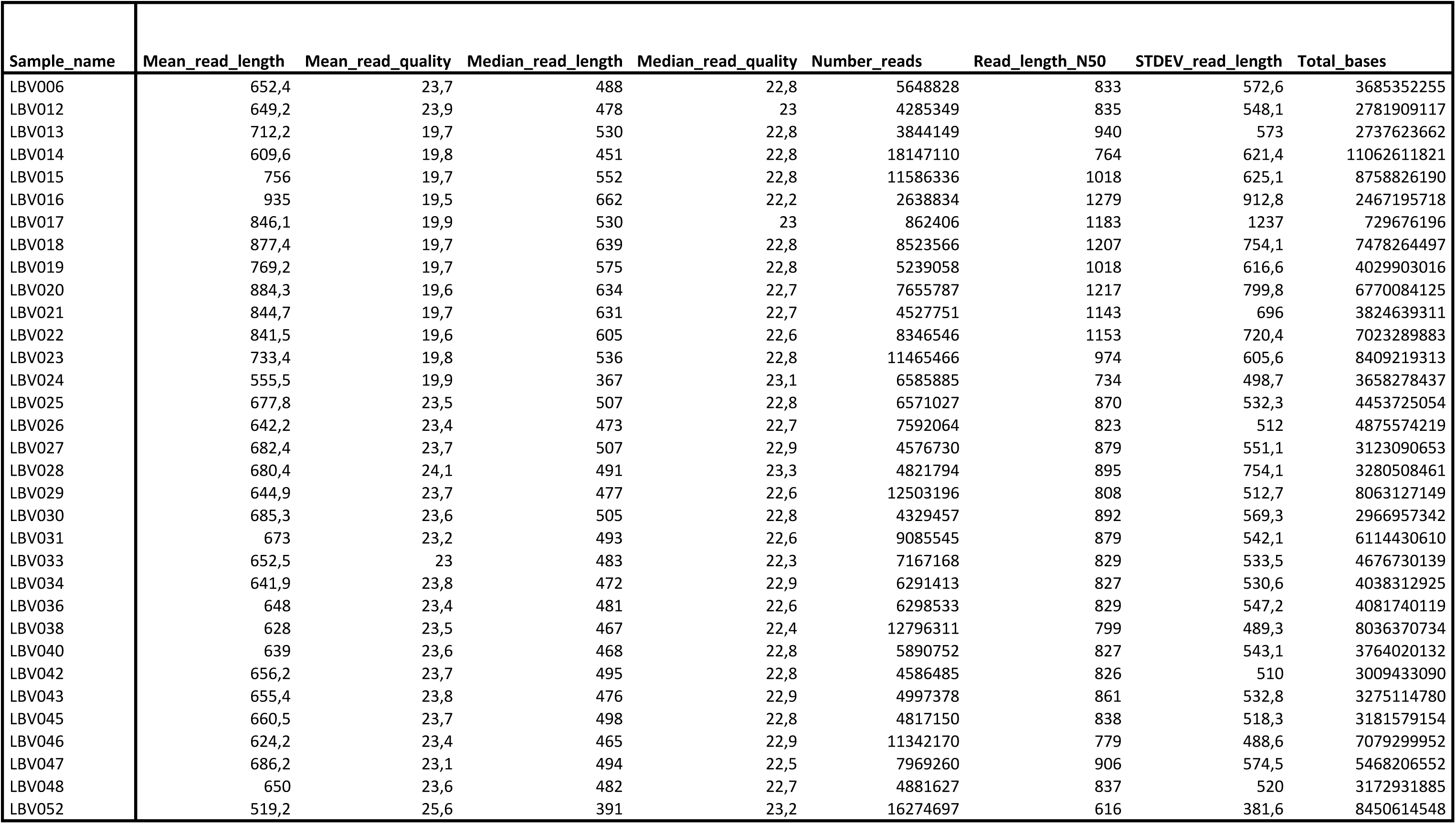
Sequencing data statistics. Summary of sequencing metrics for each isolate analyzed in the study. The table includes mean and median read lengths, read quality scores, number of reads, and total bases sequenced for each isolate.

**Supplemental Table 3.**
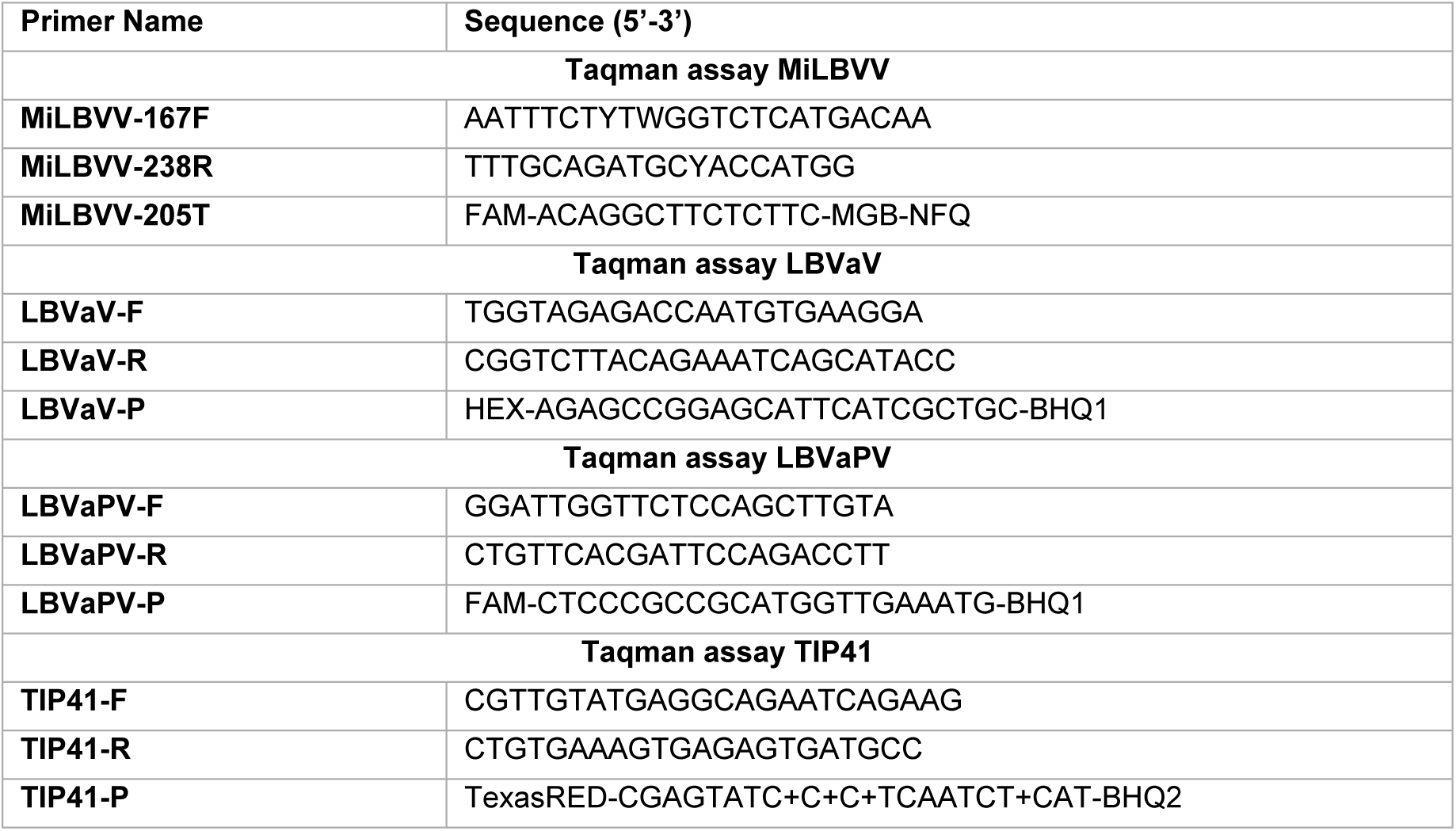
List of primers used in study. Table detailing the primers designed for RT-qPCR detection of MiLBVV, LBVaV, LBVaPV and the *L. sativa* housekeeping gene TIP41. Primer sequences are provided for forward, reverse, and probe oligonucleotides.

**Supplemental table 4.**
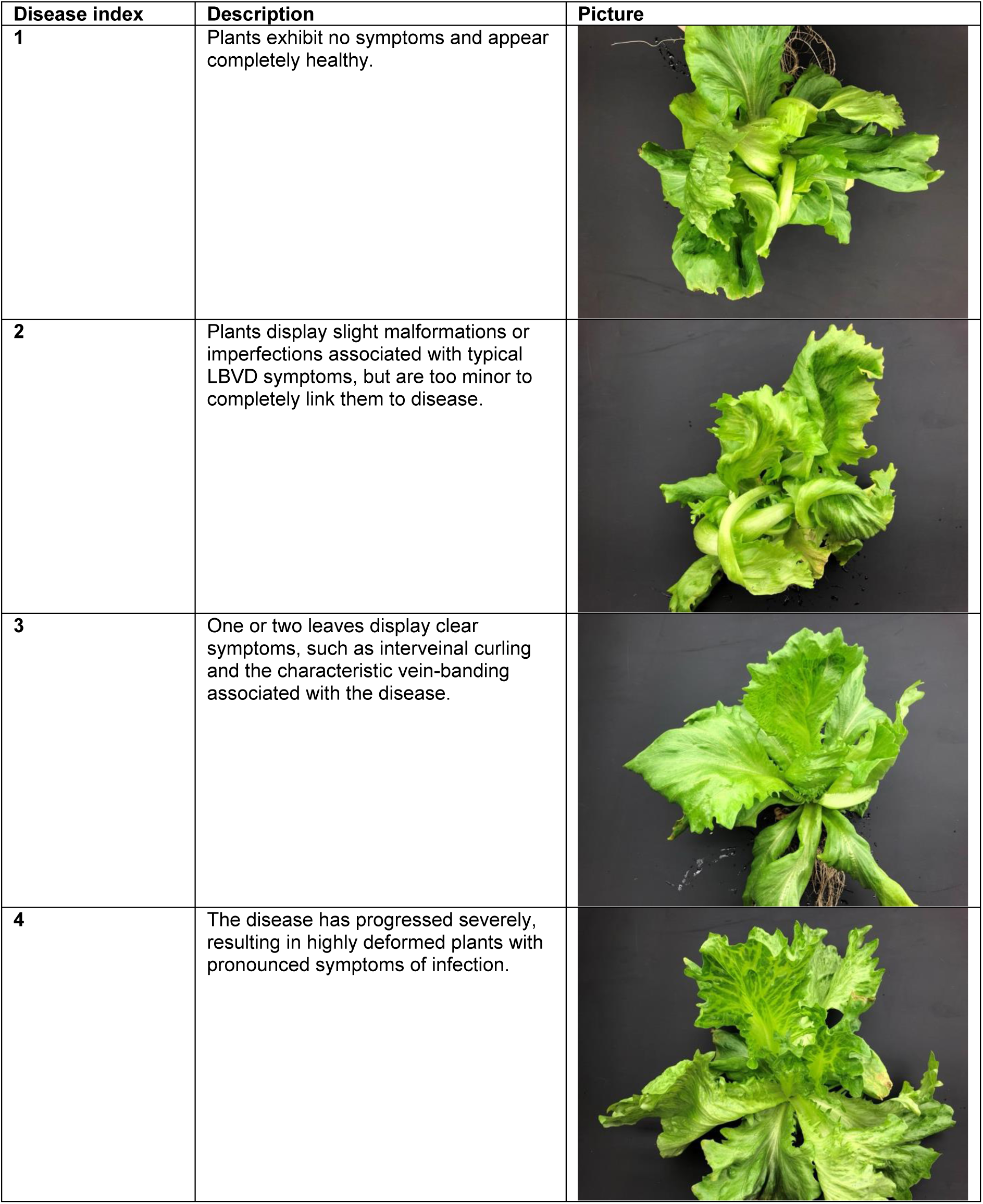
Disease index scale. Description of the disease index (DI) scoring system used to assess LBVD symptom severity in lettuce plants. The table provides a detailed description of the four DI categories, ranging from 1 (no symptoms) to 4 (severe symptoms), with representative images for each score.

| Disease index | Description | Picture |
| --- | --- | --- |
| 1 | Plants exhibit no symptoms and appear completely healthy. |  |
| 2 | Plants display slight malformations or imperfections associated with typical LBVD symptoms, but are too minor to completely link them to disease. |  |
| 3 | One or two leaves display clear symptoms, such as interveinal curling and the characteristic vein-banding associated with the disease. |  |
| 4 | The disease has progressed severely, resulting in highly deformed plants with pronounced symptoms of infection. |  |

